# The mechanism of coupled folding-upon-binding of an intrinsically disordered protein

**DOI:** 10.1101/2020.03.23.004283

**Authors:** Paul Robustelli, Stefano Piana, David E. Shaw

## Abstract

Intrinsically disordered proteins (IDPs), which in isolation do not adopt a well-defined tertiary structure but instead populate a structurally heterogeneous ensemble of interconverting states, play important roles in many biological pathways. IDPs often fold into ordered states upon binding to their physiological interaction partners (a so-called “folding-upon-binding” process), but it has proven difficult to obtain an atomic-level description of the structural mechanisms by which they do so. Here, we describe in atomic detail the folding-upon-binding mechanism of an IDP segment to its binding partner, as observed in unbiased molecular dynamics simulations. In our simulations, we observed over 70 binding and unbinding events between the α-helical molecular recognition element (α-MoRE) of the intrinsically disordered C-terminal domain of the measles virus nucleoprotein (N_TAIL_) and the X domain (XD) of the measles virus phosphoprotein complex. We found that folding-upon-binding primarily occurred through induced-folding pathways (in which intermolecular contacts form before or concurrently with the secondary structure of the disordered protein)—an observation supported by previous experiments—and that the transition state ensemble was characterized by the formation of just a few key intermolecular contacts, and was otherwise highly structurally heterogeneous. We found that when a large amount of helical content was present early in a transition path, N_TAIL_ typically unfolded, then refolded after additional intermolecular contacts formed. We also found that, among conformations with similar numbers of intermolecular contacts, those with less helical content had a higher probability of ultimately forming the native complex than conformations with more helical content, which were more likely to unbind. These observations suggest that even after intermolecular contacts have formed, disordered regions can have a kinetic advantage over folded regions in the folding-upon-binding process.

## Introduction

In isolation, intrinsically disordered proteins (IDPs) do not adopt a well-defined tertiary structure, but rather populate a heterogeneous ensemble of interconverting states. The biological interactions of IDPs are often mediated through sequence segments that can undergo disorder-to-order transitions upon interacting with structured binding partners (a so-called “folding-upon-binding” process). As IDPs populate many structurally diverse conformations when unbound, they are often able to form low-affinity but highly specific interactions with multiple binding partners—an ability that facilitates some of the important roles played by IDPs in signal transduction, cellular regulation, and various other biologically important processes.^1^

A central challenge in the study of IDPs is the characterization of the mechanisms by which they bind their physiological interaction partners: Mechanistic insight into the folding-upon-binding process of IDPs could ultimately enable a more predictive understanding of how their sequences, conformational propensities, and biophysical properties dictate their interactions and thus their biological activity. An increasing number of experimental,^2–6^ theoretical,^7–11^ and computational^12–21^ studies have been used to predict or globally characterize molecular recognition in IDPs, but atomic-resolution details have only recently begun to emerge.^3–5^

Atomistic molecular dynamics (MD) simulations are a promising approach for complementing experimental measurements of IDP folding-upon-binding. In the investigation reported here, we employed unbiased, all-atom MD simulations to study the mechanism of folding-upon-binding of the α-helical molecular recognition element (α-MoRE) of the intrinsically disordered C-terminal domain of the measles virus nucleoprotein (N_TAIL_) to the X domain (XD) of the measles virus phosphoprotein complex. Using the recently developed a99SB-*disp*^15^ force field, which provides improved descriptions of the secondary structure propensities and dimensions of disordered proteins while maintaining accurate descriptions of folded proteins, we were able to simulate the reversible folding and binding of the N_TAIL_ α-MoRE to XD near the simulated melting temperature of the complex. In these simulations, we observed over 70 binding and unbinding events, enabling us to quantify the global folding-upon-binding free-energy landscape, characterize the binding and unbinding pathways in atomic detail, and determine the transition state ensemble (TSE).

In our simulations we observed substantial heterogeneity in the binding transitions; the TSE was thus quite structurally heterogeneous, and we found that it was characterized by the formation of just a few key contacts. Folding-upon-binding primarily occurred through pathways in which intermolecular contacts formed before or concurrently with the secondary structure of the N_TAIL_ α-MoRE, a result consistent with a so-called “induced-folding” mechanism and supported by previous experimental kinetics measurements.^6,11^ Pathways in which folding of the N_TAIL_ α-MoRE preceded the formation of intermolecular contacts (consistent with a so-called “conformational selection” mechanism) played a negligible role.

We observed that the majority of transition paths proceeded through flexible states containing relatively low helical content, and large helices that formed early in transition paths generally unfolded, refolding only later in the pathway after additional intermolecular contacts were formed. We also observed that among conformations with the same number of intermolecular contacts, conformations with less helical content, and thus more conformational flexibility, had a higher probability of forming additional intermolecular contacts and ultimately forming the stable native complex than did more helical conformations. These observations suggest that early secondary structure formation does not always facilitate progress along folding-upon-binding transition paths, and that disordered regions may have a kinetic advantage over folded regions in progressing from an encounter complex to the stable bound state.^7^ Conformational disorder may thus facilitate successive contact formation during folding-upon-binding pathways, in addition to its proposed fly-casting^22^ role in facilitating encounter complex formation..

## Results

### Apo ensemble of the N_TAIL_ α-MoRE

We performed a 100 μs unbiased MD simulation of the 21-residue N_TAIL_ α-MoRE (residues 484–504) of the intrinsically disordered C-terminal domain of the measles virus nucleoprotein N_TAIL_ at 300 K using the a99SB-*disp* force field,^15^ which was developed to provide accurate descriptions of both folded and disordered protein states and, importantly, alleviates force field deficiencies that produce overly compact ensembles of disordered proteins.

We compared the helical propensity (as calculated by the STRIDE algorithm^23^) of the N_TAIL_ α-MoRE from this unbiased simulation to the helical propensity of a previously published NMR ensemble^24^ (calculated with the ASTEROIDS ensemble selection algorithm^24–26^ using NMR chemical shifts and residual dipolar couplings (RDCs) as restraints), and the results are shown in Figure 1. The simulated helical propensity is in excellent agreement with that of the experimentally derived ensemble. We also compared the agreement of both NMR chemical shifts and RDCs back-calculated from the MD simulation and chemical shifts and RDCs calculated from the ASTEROIDS ensemble with the experimental data (Table 1). The agreement with experimental chemical shifts observed in the unbiased MD simulation is comparable to the agreement observed in the ASTEROIDS ensemble, which enforced the chemical shifts as restraints, while RDCs calculated from the MD ensemble are in worse agreement with the experimental RDCs than those calculated from the ASTEROIDS ensemble (Figure S2), particularly for residues R489, R490, and S491. As these residues are located near the N-terminal region of the N_TAIL_ α-MoRE and protrude into the solvent in the N_TAIL_:XD complex structure, we do not expect this relatively minor discrepancy in the apo ensemble to have a large impact on the simulated folding-upon-binding mechanism.

**Figure 1.**
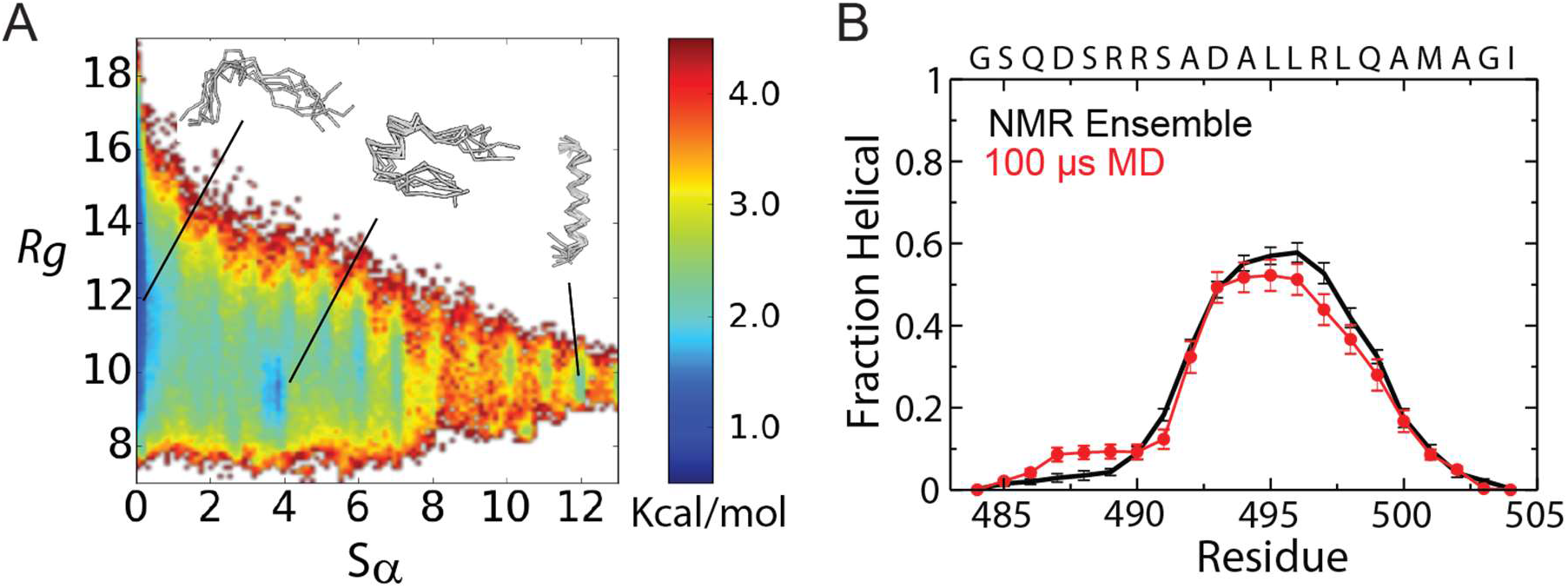
Ensemble of the apo N_TAIL_ α-MoRE from a 100-μs MD simulation at 300 K. A) Free energy surface of the apo N_TAIL_ α-MoRE as a function of the radius of gyration (R_g_) and the α-helical folding order parameter Sα. B) Comparison of the helical propensity of the apo N_TAIL_ α-MoRE calculated from a 100-μs MD simulation and from an NMR ensemble^24^ calculated with the ASTEROIDS ensemble selection^25^ algorithm using NMR chemical shifts and RDCs as restraints. Helical states were assigned using the program STRIDE.^23^

**Table 1.**
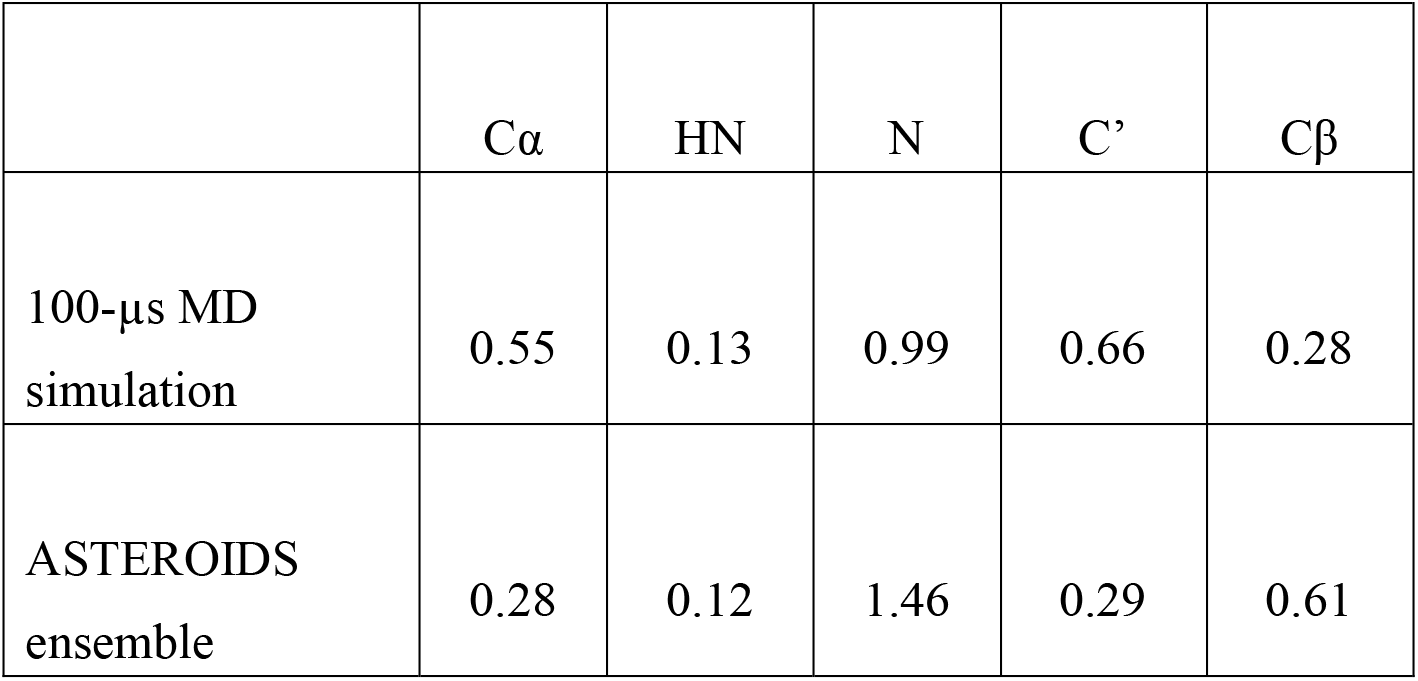
RMSD between calculated and experimental NMR chemical shifts (ppm) from a 100 μs unbiased MD simulation using the a99SB-*disp* force field and from an NMR ensemble^24^ calculated with the ASTEROIDS ensemble selection^25^ algorithm, using NMR chemical shifts and RDCs as restraints. Chemical shifts were calculated using SPARTA+.^47^ Calculated RMSDs for both ensembles are less than the reported error of SPARTA+ predictions on databases of folded X-ray structures.

| | Cα | HN | N | C' | Cβ |
| --- | --- | --- | --- | --- | --- |
| 100-μs MD simulation | 0.55 | 0.13 | 0.99 | 0.66 | 0.28 |
| ASTEROIDS ensemble | 0.28 | 0.12 | 1.46 | 0.29 | 0.61 |

We visualize the conformational free-energy surface of the apo N_TAIL_ α-MoRE as a function of the radius of gyration (Rg) and the α-helical folding order parameter Sα^27^ (Figure 1). Sα is a measure of α-helical content that reports the similarity of each consecutive 6-residue segment of a protein to the conformation of an ideal α-helix (See Methods). A 6-residue segment that has a backbone RMSD from an ideal α-helical conformation of <0.4 Å has an Sα value very close to 1, and a segment with a backbone RMSD from an ideal α-helical conformation of >4.0 Å has an Sα value very close to 0. A fully helical conformation of the N_TAIL_ α-MoRE has an Sα value of 13, and a conformation of the N_TAIL_ α-MoRE with no helical content has an Sα value of 0.

The most populated minimum in the MD ensemble was a disordered state with an Sα value <1 (39.8% of the ensemble). There was a relatively low population of states with an Sα value >7 (5.6%), and a complete helix—with an Sα value greater than 12.5—rarely formed in the apo ensemble (0.3% of the ensemble). There was a relatively compact free-energy minimum centered between 3 < Sα < 4, in which the N-terminal and C-terminal tails interacted behind a central helical turn (13.0% of the ensemble). We note that the N_TAIL_ α-MoRE MD ensemble calculated here with the a99SB-*disp* force field is substantially more expanded than previously reported N_TAIL_ α-MoRE MD ensembles^12,14^ calculated with the CHARMM22* force field^28^ (average R_g_ = 11.0 Å with a99SB-*disp* compared to ~8–8.5 Å with CHARMM22*) and shows improved agreement with secondary Cα chemical shifts (r = 0.92) compared to the agreement previously reported for a CHARMM22* MD ensemble (r = 0.76).^14^

### Simulations of folding-upon-binding of the N_TAIL_ α-MoRE to XD

In an effort to observe a large number of binding and unbinding transitions in an equilibrium simulation, we sought to simulate the association of the N_TAIL_ α-MoRE to XD at a temperature near the simulated melting temperature of the complex. We found that in simulations run at 400 K the α-MoRE:XD complex was bound in 74% of the simulation, the XD domain remained folded in the unbound state, and we were able to frequently observe binding and unbinding transitions (Figure 2). We monitored complex formation using the fraction of intermolecular contacts *Q*, and defined binding and unbinding events as transitions between states where *Q* = 0 and *Q* > 0.95, using a dual-cutoff approach.^29^ In a 200-μs simulation we observed 36 binding and 36 unbinding transitions.

**Figure 2.**
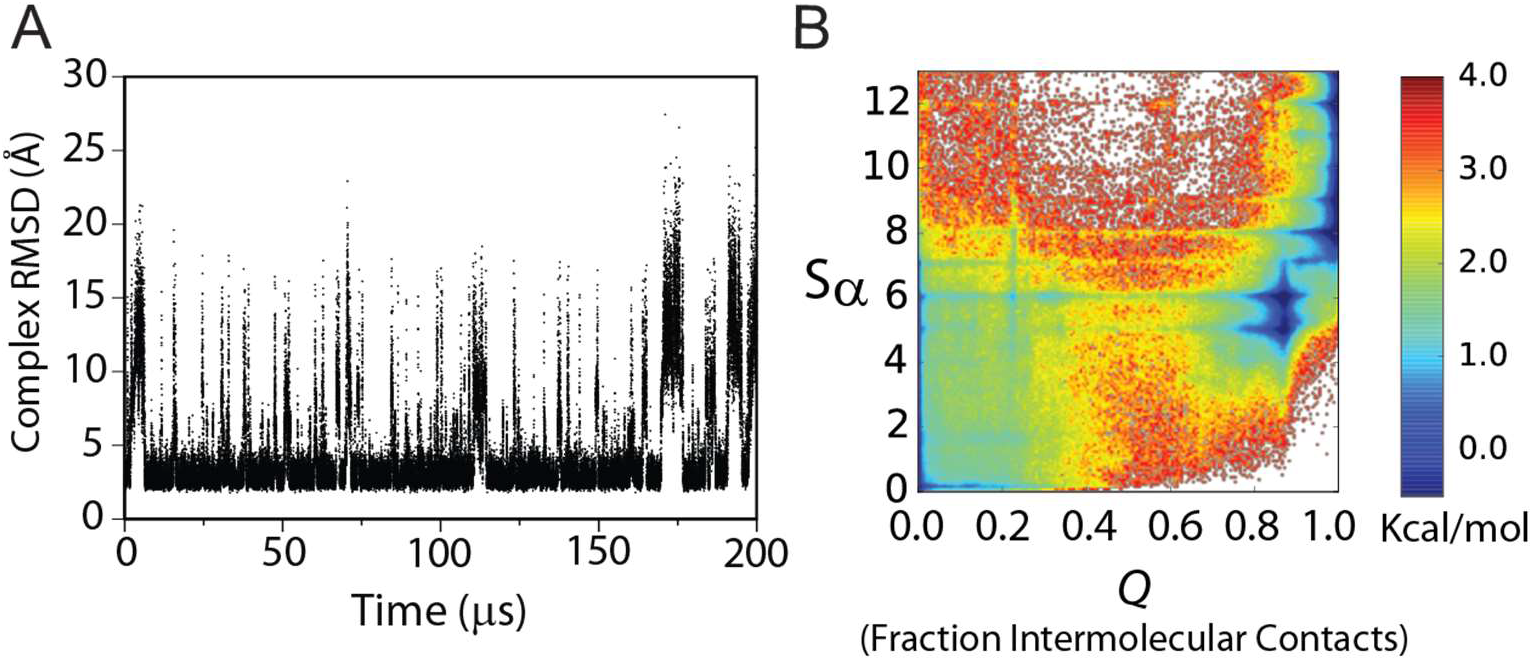
A) RMSD of the N_TAIL_ α-MoRE and XD from the native N_TAIL_ α-MoRE:XD complex (PDB 1T6O) in an unbiased 200-μs MD simulation run with the a99SB-*disp* force field at 400 K. B) Free energy surface of complex formation as a function of the N_TAIL_ α-MoRE α-helical folding order parameter Sα and the fraction of native intermolecular contacts Q.

### Free energy surface of folding-upon-binding of the N_TAIL_ α-MoRE to XD

We visualize the free-energy surface of complex formation as a function of *Q* and Sα in Figure 2. Analysis of the free-energy surface (Figure 2) and of intermolecular contact formation during N_TAIL_ α-MoRE folding-upon-binding transitions paths (Figure 3) indicates that there is a very low probability of observing pure conformational selection events, in which a completely helical N_TAIL_ α-MoRE binds to XD, or pure induced folding events, in which a completely disordered N_TAIL_ α-MoRE forms all intermolecular contacts before folding into a helix. Instead, the free-energy surface and transition path analysis are consistent with a scenario in which both the formation of intermolecular contacts and the folding of the N_TAIL_ α-MoRE occur concurrently. The encounter complexes observed in simulation (defined as 0 < *Q* < 0.1) had, on average, a relatively low helical content (Sα = 3.7), not much higher than the isolated N_TAIL_ α-MoRE (Sα = 2.7), indicating that the folding-upon-binding pathways more closely resemble an induced-folding mechanism than a conformational-selection mechanism. A free-energy minimum, corresponding to a partially helical intermediate state that has formed the majority of intermolecular contacts, is observed at *Q* = 0.85 and Sα = 5. States with more helical content than this intermediate state were only substantially populated when almost all of the native contacts had already formed (*Q* > 0.95), suggesting that progression from this partially folded helical state to the full helix occurred through a pure induced folding mechanism.

**Figure 3.**
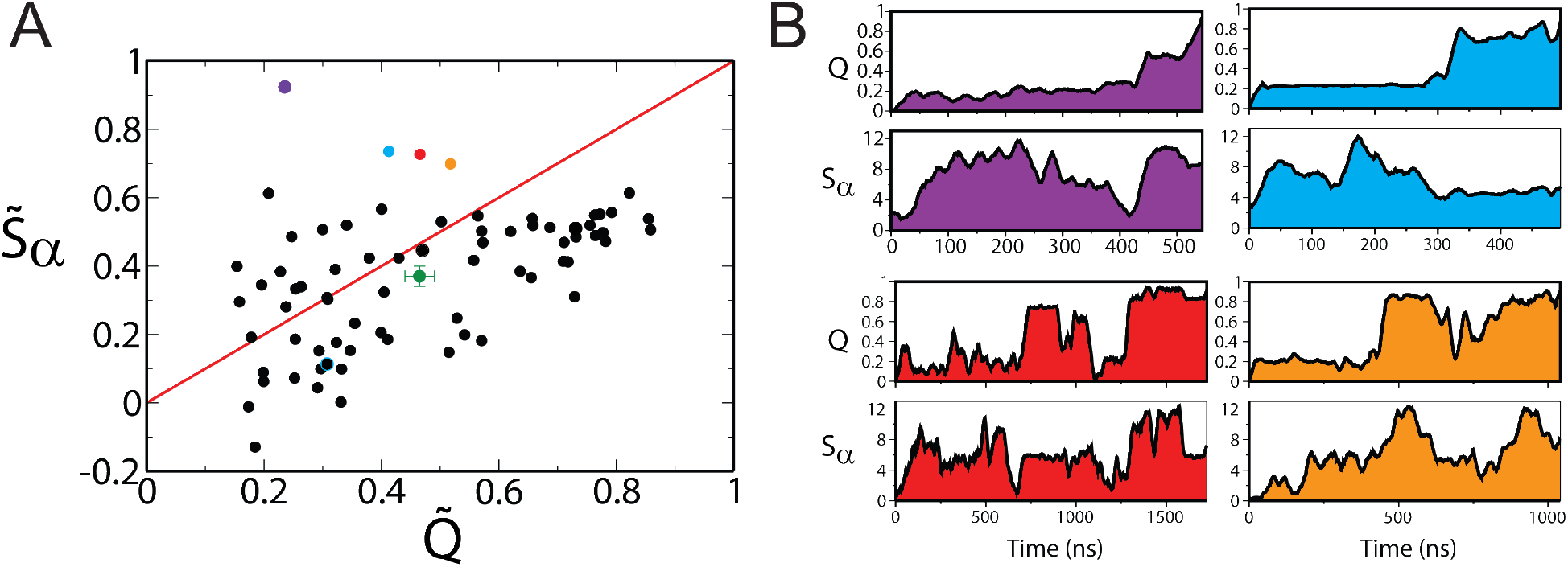
A) Comparison of 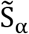 and 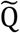 (normalized time averages of Sα and Q) for 72 binding and unbinding transition paths observed in an unbiased 200-μs MD simulation of the N_TAIL_ α-MoRE and XD domain run with the a99SB-*disp* force field at 400 K. 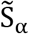 and 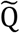 are calculated by integrating Sα and Q for a given transition path and normalizing such that values of 0 and 1 correspond to the average values of Sα and Q in the unbound and bound states, respectively (see Methods). The average value of 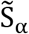 and 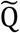 over all transition paths is shown in green. The four transition paths with the largest values of 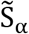 are indicated by the purple, blue, red, and orange circles. B) The time-evolution of Sα and Q for the four transition paths with the largest values of 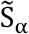. We note that the helical structure formed early in these transition paths unfolds before additional intermolecular contacts are formed.

### Analysis of helix and contact formation during the folding-upon-binding transition paths

To obtain more direct insight into the folding-upon-binding mechanism, we analyzed the formation of intermolecular contacts and N_TAIL_ α-MoRE folding in the 72 binding and unbinding transition paths. Unbinding events were analyzed in reverse, such that all transitions could be treated as binding events. 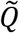 and 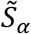 (normalized time averages of *Q* and Sα) were calculated by integrating *Q* and Sα for a given transition path and normalizing such that values of 0 and 1 correspond to the average values of *Q* and Sα in the unbound and bound states, respectively (see Methods). In Figure 3 we plot 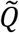 and 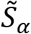 for each transition path, and illustrate the time evolution of *Q* and Sα for the four transition paths with the largest values of 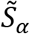. 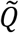 and 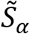 provide information about the folding-upon-binding mechanism, as they reflect how early in a transition path the formation of intermolecular contacts and helix folding occurred, with high values reflecting early formation.^29^ The 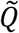 and 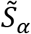 of mechanisms in which native contacts and helical turns form simultaneously during the transition would fall along the diagonal (red line in Figure 3).

We find that 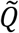 and 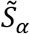 are roughly scattered along the diagonal, but 71% of transition paths have a larger 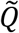 than 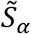. These results indicate that the formation of intermolecular contacts and N_TAIL_ α-MoRE folding occurred relatively concurrently, but that, on average, the formation of intermolecular contacts occurred earlier in transition paths than an increase in the helical content of the N_TAIL_ α-MoRE relative to the apo state (a result consistent with a mechanism characterized by induced folding more than conformational selection). We examined if the observed mechanisms of folding-upon-binding change with temperature by running 25 unbiased binding simulations at 300 K, in which an unfolded conformation of the N_TAIL_ α-MoRE was placed in a water box containing the XD domain and the simulation was run until a stable complex formed. We found that the distribution of folding-upon-mechanisms in these 22 binding events was similar to that observed at 400 K (Figures S5 and S6), but that the transition paths tended have even more of an induced-folding character at 300 K.

While the majority of the folding-upon-binding transition paths observed in simulations (at both 400 K and 300 K) had an induced-folding character, three transition paths were more consistent with a conformational-selection mechanism, as their average helical content was >80% of the difference between bound-state helicity and unbound helicity. In five cases, the transition paths featured conformations that, on average, had *less* helical content than the N_TAIL_ α-MoRE did in apo simulations (i.e., 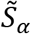 is <0). In these instances, after forming initial contacts, the binding pathway predominantly proceeded through states that were less helical than the apo N_TAIL_ α-MoRE, illustrating a scenario in which unfolding, relative to the apo form, facilitated the formation of the stable native complex. We thus observed more transition paths in which the N_TAIL_ α-MoRE exhibited what we term an *unfolding-to-bind* mechanism, in which binding transitions of a disordered protein predominantly proceed through conformations that have less helical content than its apo form, than transition paths that exhibited a conformational-selection mechanism.

Examination of the time-evolution of *Q* and Sα provide additional insight into the nature of the binding and unbinding events. In Figure 3B we plot the time evolution of *Q* and Sα for the 4 transition paths at 400 K with largest integrals of Sα. In each of these transition paths, long helices formed early in the pathway. These helices, however, did not remain stable throughout the binding event, but rather unfolded before additional intermolecular contacts formed, and only refolded later in the path, concurrently with intermolecular contact formation. This observation is in contrast to the conformational-selection paradigm, in which a stable, preformed helix remains largely formed for the duration of the transition path. Furthermore, we also observed that in transition paths that had more induced-folding character (Figure S4), previously formed intermolecular contacts sometimes broke before the folding of additional helical elements, and then reformed. These results suggest that the barrier for folding-upon-binding transition is broad and relatively flat, such that both contact formation and helix formation are somewhat diffusive processes; there is not a completely clear ordering or separation of timescales for contact formation and helix formation (a finding consistent with conclusions from previous simulation studies^20,21^).

To further quantify the level of correlation between helix and contact formation, we discretized the trajectories into 50-ns windows and calculated the variations in the average Sα and *Q* values for each window. We observed that increases in Sα between consecutive windows were accompanied by increases in *Q* 66% of the time, and decreases in *Q* 24% of the time (with *Q* remaining constant within an interval of 0.01 the remaining 11 % of the time). We observed that increases in *Q* were accompanied by increases in Sα 62 % of time and decreases in Sα 26 % of the time (with Sα remaining constant within an interval of 0.10 the remaining 12 % of the time) (Table S1). We report the distribution of the correlation coefficients for the values of Sα vs. *Q* for all 72 transition paths in Figure S3. The correlation coefficient of Sα vs. *Q* taken over all transition paths is 0.55.

Finally, to better understand the role of helix formation in folding-upon-binding pathways, we performed a reactive flux, or P_Fold_, analysis of the entire 200 μs trajectory (Figure 4).^30^ We coarse-grained the trajectory into microstates using even intervals of Sα and *Q*, partitioning the trajectory into rectangular grid cells. For each microstate, we calculated the probability that conformations from that microstate would successfully fold before unbinding (P_Fold_). We observed that, at low to intermediate *Q* values (0.2 < *Q* < 0.6), while conformations with some secondary structure had higher P_Fold_ values than completely disordered conformations, conformations with high helical content (Sα > 6) were less likely to successfully fold and bind than less-helical conformations (with 4 < Sα < 6), suggesting that states with intermediate levels of helicity and more conformational flexibility are more likely to make additional intermolecular contacts and ultimately form the stable native complex before dissociating than more helical states. These observations, together with our analyses of the integrals of Sα and *Q* observed in transition paths, suggest that in the case of the N_TAIL_ α-MoRE binding the XD domain, early formation of large amounts of secondary structure does not significantly facilitate progress along folding-upon-binding transition paths, in contrast to the classical notion of conformational selection.

**Figure 4.**
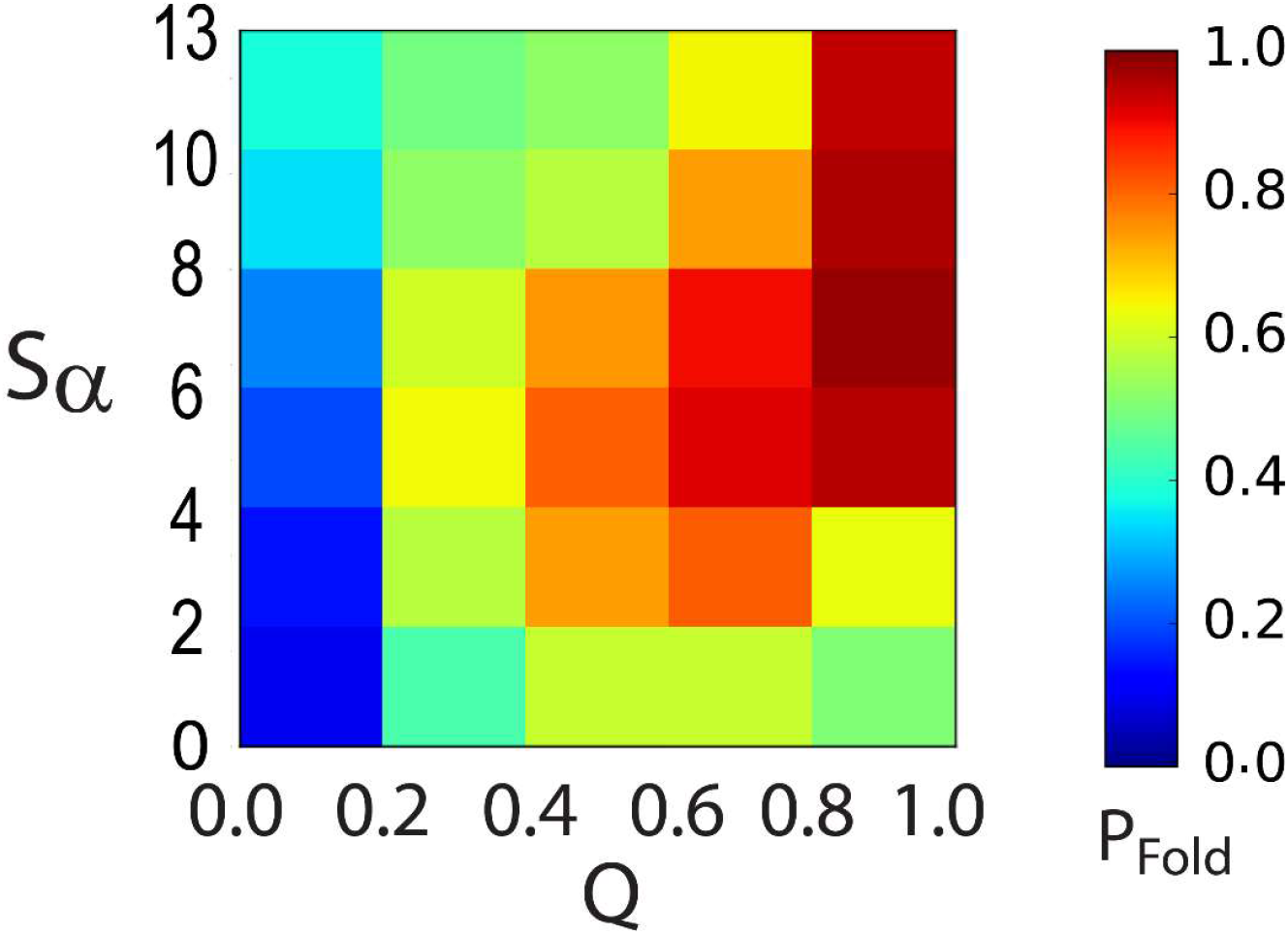
The probability of the N_TAIL_ α-MoRE folding and binding to XD (P_fold_) as a function of the α-helical folding order parameter Sα and the fraction of native intermolecular contacts *Q* calculated from an unbiased 200-μs MD simulation run with the a99SB-*disp* force field at 400 K.

### Optimization of Folding-Upon-Binding Reaction Coordinate

To further characterize the molecular mechanism and the transition state of folding-upon-binding, we next sought to optimize an improved reaction coordinate (relative to the fraction of native intermolecular contacts *Q*) that better describes the progress of folding-upon-binding reactions and defines a meaningful TSE. Starting with the reaction coordinate *Q*, we used the variational optimization algorithm proposed by Best and Hummer^31^ to reweight the contribution of each intermolecular contact to optimize a new reaction coordinate, *R* (see Methods). The results of the optimization are shown in Figure 5. Although there is some correlation in the data, and different weights can be obtained using different optimization parameters, the main features of the optimized coordinate are rather robust: The new reaction coordinate *R* reduced the weights of most contacts to values close to 0, while leaving 4 contacts with values near 1. 1D and 2D projections of the free energy landscape of the simulation as a function of *R* and of *R* and Sα, respectively, are shown in Figure 6. 2D projections of the probability of being on a transition path as functions of *Q* and Sα (p(TP|*Q*,Sα)) and of *R* and Sα (p(TP|*R*,Sα) are shown in Figure S7.

**Figure 5.**
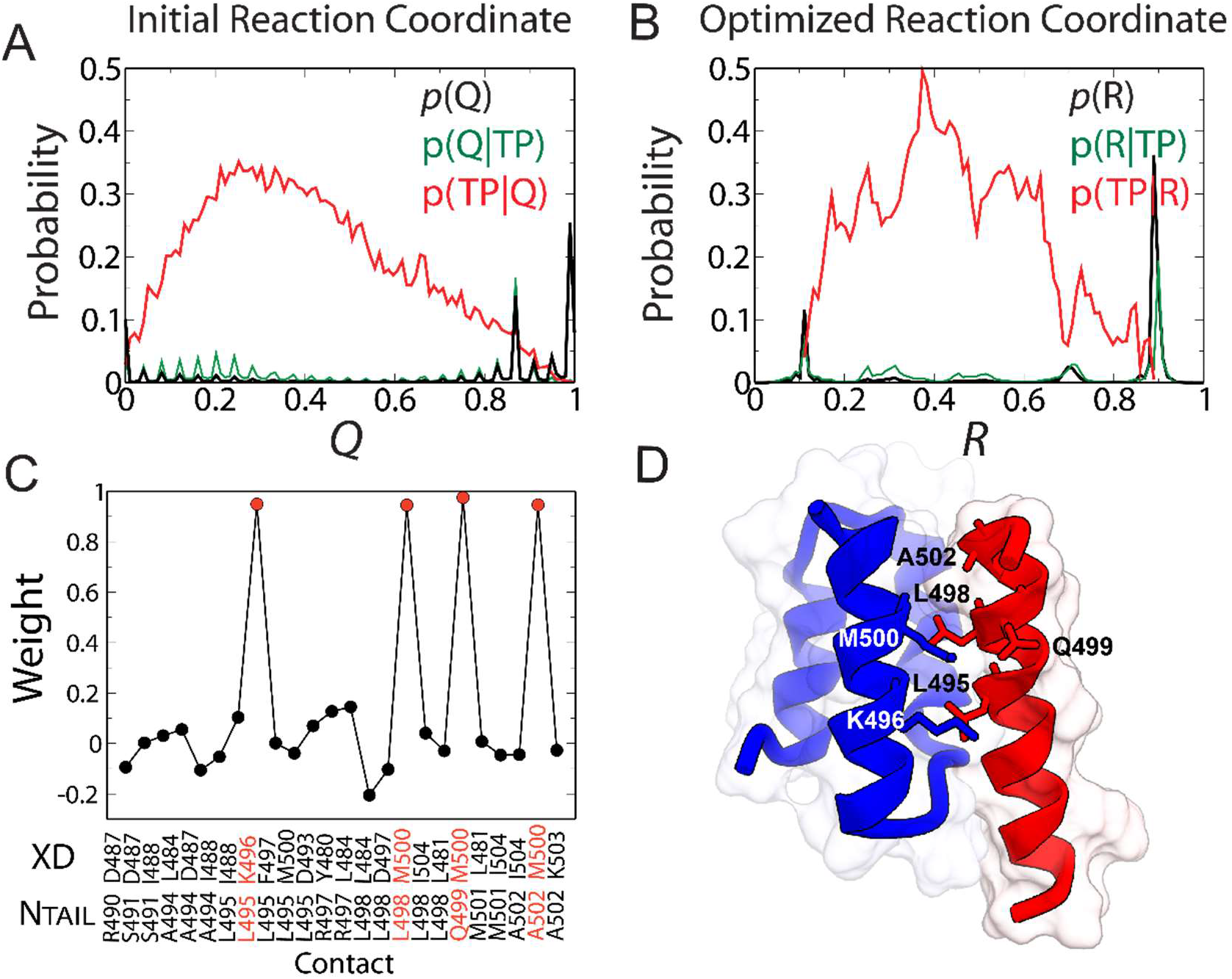
Optimization of a new reaction coordinate *R* to better describe folding-upon-binding. A) The equilibrium probability of the original reaction coordinate *Q* (p(*Q*), black), the probability of *Q* given that a frame is on a transition path (p(*Q*|TP), green), and the conditional probability of being on a transition path given a value of *Q* (p(TP|*Q*), red). B) The corresponding probabilities for the optimized reaction coordinate *R*. C) The weights of each intermolecular contact in the new reaction coordinate *R*. D) The structure of the N_TAIL_ α-MoRE:XD complex. Residues forming contacts with the largest weights in the optimized reaction coordinate *R* are shown as sticks and labeled.

**Figure 6.**
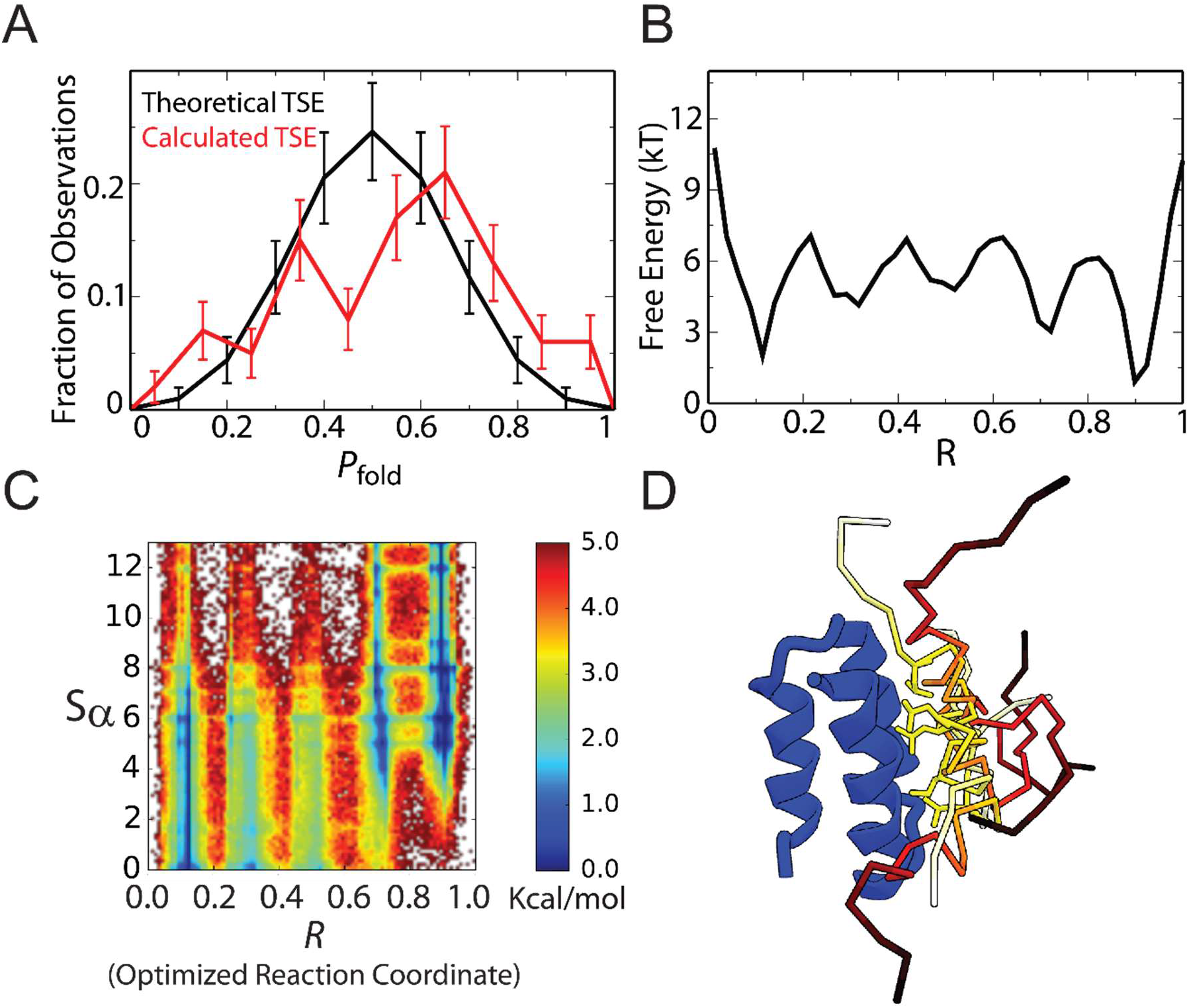
Calculation of a TSE for folding-upon-binding. A) Distribution of the probability of folding and binding (P_fold_) for 100 frames from a 200-μs MD a99SB-*disp* trajectory; the frames were selected randomly from frames with 0.3 < *R* < 0.5 (such frames are predicted to have a high probability of being on a transition path). B) Free energy of the trajectory as a function of *R*. C) The free energy of complex formation as a function of the N_TAIL_ α-MoRE α-helical folding order parameter Sα and *R*. D) Five representative structures of the TSE of folding-upon-binding. Residue L498 is displayed in stick representation (yellow).

### Calculation of a transition state ensemble

To test the quality of the new reaction coordinate *R*, we randomly selected 100 frames from the simulation with values of *R* that are predicted to have the highest probability of being on a transition path (0.3 < *R* < 0.5). 50 frames were selected that had previously been assigned to transition paths and 50 frames were selected that were not previously assigned to transition paths. For each frame, we ran 10 “shooting” simulations, in which atoms were assigned random velocities, and simulations were run until a dissociation event (*Q* = 0) or a binding event (*Q* > 0.95) was observed. For each starting structure, we computed the probability of folding and binding (P_fold_) from the shooting simulations. In Figure 6A, the observed distribution of P_fold_ is compared to the expected binomial distribution for 10 observations if all structures were part of a TSE and had a P_fold_ value of exactly 0.5. The distribution is in reasonable agreement with the theoretical distribution, suggesting that *R* is a good reaction coordinate for folding-upon-binding, with a slight bias toward the bound state.

We defined a final TSE by selecting all tested frames with P_fold_ values between 0.3 and 0.7. A subset of these structures is shown in Figure 6C. We found that the TSE was notable in that there does not appear to be a preferred orientation of the N_TAIL_ α-MoRE relative to the XD domain. The average helicity of each residue and the contact profile of the TSE is shown in Figure S8. In the TSE the relative helicity of each residue was very similar to its helicity in the apo ensemble of the N_TAIL_ α-MoRE at 400 K, although, overall, the TSE was slightly more helical. There is substantial variability in the distribution of the location of helices within the individual conformations of the TSE. The most frequently formed intermolecular contacts in the TSE involved residues L498 and L495, and the most populated contact is between L498 of the N_TAIL_ α-MoRE and M500 of XD (formed in 65% of the TSE conformations). The unifying feature of the TSE appears to be the insertion of L498 and L495 in the binding groove of XD, while the structure and relative orientation of the N_TAIL_ α-MoRE are highly heterogeneous.

## Discussion

We report here a fully atomistic description of the folding-upon-binding of the N_TAIL_ α-MoRE to the XD domain as obtained by unbiased equilibrium MD simulations performed near the melting temperature of the complex using a state-of-the-art physics-based force field that overcomes recently documented deficiencies in descriptions of the secondary structure propensities and dimensions of disordered proteins. The reasonably large number of folding and unfolding events observed in our simulation ensures that the observations made in this study are statistically meaningful, which is of particular importance given that the mechanism of formation of this complex appears to be highly heterogeneous.

The results of our unbiased MD simulations are largely consistent with previously reported experiments. The complex binding kinetics observed in simulation echoes findings from NMR chemical shift titration experiments, which produced results inconsistent with a simpler two-state binding process.^32^ The dominance of pathways with more induced-folding character, in which folding occurs late in the transition path, is consistent with kinetics measurements previously performed on this system, which could clearly resolve separate rates for the formation of an initial encounter complex and subsequent folding.^6^ The intermediate partially folded state observed in our simulations, which has some helical content but remains highly dynamic, is broadly consistent with relaxation dispersion NMR measurements of the homologous Sendai N_TAIL_ and XD domains, which provided a detailed characterization of an intermediate dynamic encounter complex. The encounter complex was found to contain an elevated population of helix relative to apo N_TAIL_, as assessed by backbone carbon chemical shifts, but remained dynamic and relatively nonspecifically bound, as assessed by relatively small changes in nitrogen and proton backbone chemical shifts.^3^

Additional kinetics measurements performed on N_TAIL_ mutants have previously been used to further characterize the folding and binding steps in N_TAIL_:XD complex formation.^11^ Phi-values determined for the binding step indicated that encounter complex formation is mediated by residues in the central helix of N_TAIL_, and is predominantly driven by hydrophobic interactions involving residues A494, L495, L498, and A502. Our simulations are also consistent with these observations: Optimization of the reaction coordinate *R* indicates that the intermolecular contacts formed by L495, L498, and A502 are three out of the four most important contacts for describing the folding-upon-binding process, and we observed that the insertion of residues of L495 and L498 into the XD biding groove is the predominant unifying feature of conformations of the TSE.

In our simulations, the transition state for folding-upon-binding appears to be determined by the formation of just a few key native contacts. These requirements do not impose a large number of constraints on the remaining structure and, as a result, the transition state ensemble is structurally extremely diverse, featuring helical and non-helical structures and a range of different orientations of the peptide backbone chain. Our calculated TSE is thus similar to a recently described highly heterogenous transition state for the NCBD:ACTR complex calculated using phi-value restraints.^33^

Our results are inconsistent with a conformational selection mechanism, as they indicate that the folding transition proceeds through the formation of hydrophobic contacts, with fully helical states only observed after the transition state, when the majority of the native contacts have already been formed. In fact, our simulations suggest the existence of an “unfolding-to-bind” mechanism, in which disorder facilitates formation of additional key contacts; as a result, highly structured states encountered before the transition state often unfold to make further progress towards complex formation. These observations suggest that, in at least some instances, disordered regions may have a kinetic advantage over folded regions in progressing from an encounter complex to the stable bound state.^7^ Similarly to the proposed role of disorder in the fly-casting^22^ model of encounter complex formation (in which the larger capture radius of disordered proteins enables them to form key contacts for protein-protein recognition faster than fully structured proteins can), and to the so-called “cracking” process that facilitates allosteric conformational transitions,^34^ here conformational disorder may facilitate the formation of native contacts during the folding-upon-binding transition.

This study indicates that atomistic MD simulations performed with a state-of-the-art force field are able to provide an accurate description of the folding-upon-binding process that complements and helps in interpreting the results of experimental measurements. Future simulation studies would be particularly valuable for other examples of the many systems with complex mechanisms that do not strictly conform to the archetypes of conformational selection or induced folding.

## Methods

### MD simulation setup

MD simulations were performed using the a99SB-*disp* force field, CHARMM22 ions,^35^ and a99SB-*disp* water.^15^ Systems were initially equilibrated at 300 K and 1 bar for 1 ns using the Desmond software.^36^ Production runs were performed in the NPT ensemble^37–40^ with Anton specialized hardware^41^ using a 2.5-fs time step and a 1:2 RESPA scheme.^42^ Bonds involving hydrogen atoms were restrained to their equilibrium lengths using a version^43^ of the M-SHAKE algorithm.^44^ The Gaussian split Ewald method^45^ with a 32 × 32 × 32 mesh was used to account for the long-range part of the electrostatic interactions.

Simulations of the N_TAIL_ α-MoRE were performed in a 55 Å water box containing 5190 water molecules, 15 Cl^−^ ions, and 14 Na^+^ ions, and run for 100 μs. Simulations of the N_TAIL_ α-MoRE and the XD domain were performed in a 72 Å water box containing 10569 water molecules, 24Cl^-^ ions, and 19 Na^+^ ions. A 400 K simulation of the N_TAIL_ α-MoRE and the XD domain was initiated from PDB 1T6O^46^ and run for 200 μs. 300 K simulations of the N_TAIL_ α-MoRE and the XD were initiated with the N_TAIL_ α-MoRE in an unbound conformation and were run until a stable binding event was observed.

### MD simulation analysis

NMR chemical shifts were calculated from the apo N_TAIL_ α-MoRE simulation using SPARTA+^47^ RDCs were calculated with PALES^48^ with a local alignment of 15 residues. Helical propensities were calculated using secondary structure assignments calculated with the program STRIDE.^23^

In simulations of the N_TAIL_ α-MoRE and the XD domain, complex formation was monitored using the fraction of native intermolecular contacts *Q*,^29^ where contacts were calculated as a function of the minimum distance between atoms in two residues according to the following switching function:

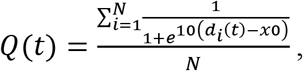

where the sum is over all *N* pairs of native intermolecular contacts between the N_TAIL_ α-MoRE and the XD domain, *d_i_* is the minimum distance between any two atoms in a pair of residues for contact *i* at time *t*. The contact cutoff was set to *x_0_* = 5 Å, such that a contact is counted as fully formed if any two atoms in a pair of residues are within 5 Å. Native intermolecular contacts were defined as those residue-residue contacts that are within 5 Å in crystal structure PDB 1T6O. Binding and unbinding events were determined as transitions between states where *Q* = 0 and Q > 0.95, using a dual-cutoff approach,^29^ where values of *Q* were smoothed using a running average with a 2 ns window.

The α-helical order parameter Sα, which measures the similarity of all 6-residue segments to an ideal helical structure, was calculated as defined by Pietrucci and Liao:^27^

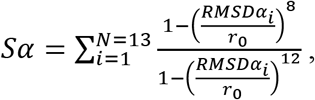

where the sum is over all *N* = 13 consecutive 6-residue segments in the N_TAIL_ α-MoRE, RMSDα_i_ is the RMSD of the backbone atoms of the 6-residue fragment *i* from an ideal α-helical geometry, and r_0_ = 0.8 Å. When r_0_ = 0.8Å, a 6-residue fragment with a value of RMSD_α_ < 0.4Å contributes a value very close to 1 to the Sα sum, and a segment with a value of RMSD_α_ > 4.0Å contributes a value very close to 0 to the Sα sum. A completely helical conformation of the N_TAIL_ α-MoRE has an Sα value of 13, and a completely disordered conformation of the N_TAIL_ α-MoRE has an Sα value of 0. In all plots values of Sα were smoothed using a running average with a 2 ns window.^29^

The probability of folding (P_fold_) as a function of *Q* and Sα was calculated by initially assigning every frame of the 400 K trajectory to a bin on an evenly spaced 6 × 6 grid according to its value of Sα and Q (Figure 4). For each bin, the P_fold_ was computed as the probability that the trajectory started from one of the frames of that bin would reach values of *Q* > 0.95 before dissociating (*Q* = 0.0).

### Optimization of a folding-upon-binding reaction coordinate

We optimized a new reaction coordinate *R* using the variational optimization algorithm proposed by Best and Hummer.^31^ We started the optimization using the reaction coordinate *Q*, where all native intermolecular contacts contribute equally to the reaction coordinate. Each frame in the trajectory was assigned as either belonging to a transition path or not belonging to a transition path. Based on this assignment, we calculated the probability distribution of *Q* for the entire trajectory (p(Q)), as well as the probability distribution of *Q* observed for all frames that were assigned to a transition path *Q* (p(Q|TP)). From these values we calculated the probability of being on a transition path for a given value of *Q* (p(TP|Q)) from the equality p(Q|TP)p(TP) = p(TP|Q)p(Q) (where p(TP) is the fraction of time spent in transition paths). For an ideal reaction coordinate, p(TP|Q) would resemble a Gaussian distribution with a maximum value of 0.5.

To optimize a new reaction coordinate *R*, we used a Monte Carlo algorithm that proposed modifications to the weight of each intermolecular contact in the definition of *R*. The starting reaction coordinate was thus 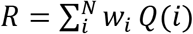, where the value of *Q*(*i*) is defined for the *i*th intermolecular contact as defined in Equation 1, and *w_i_* = 1 for all values. For each Monte Carlo move, an intermolecular contact was selected at random, and a proposed modification to the weight *w_i_* was drawn from a Gaussian distribution centered at 0 with a standard deviation of 0.2. Based on the new proposed reaction coordinate *R*’, the probability of being on a transition path for a given value of *R*’ ((p(TP|R’)) was calculated and was fit to a Gaussian function, and the score of the new reaction coordinate *R*’_score_ was assigned as the maximum value of the best-fit Gaussian function. If the new reaction coordinate *R*’ resulted in an improved score relative to the previous reaction coordinate *R*, the Monte Carlo move was accepted, and otherwise the move was accepted with an acceptance probability proportional to exp(*R*’_score_ – *R*_score_) / T, where T is a temperature. Multiple annealing schedules of *T* were tested, and the final reaction coordinate with the highest value of *R*_score_ was selected as the final reaction coordinate *R*.

### Calculation of a transition state ensemble

To calculate a TSE we randomly selected 100 frames from the 400 K simulation of the N_TAIL_ α-MoRE and the XD domain with values of *R* that were predicted to have the highest probability of being on a transition path (0.3 < *R* < 0.5). 50 frames were selected that had previously been assigned to transition paths and 50 frames were selected that were not previously assigned to transition paths. For each frame, we ran 10 shooting simulations, in which atoms were assigned random velocities, and simulations were run until a dissociation event (*Q* = 0) or a binding event (*Q* > 0.95) was observed. For each starting structure, we computed the probability of folding and binding (P_fold_) from the shooting simulations. All frames with P_fold_ values between 0.3 and 0.7 were assigned to the TSE.

## Acknowledgments

We thank Mike Eastwood for critical readings of the manuscript and providing helpful suggestions, and Berkman Frank for editorial assistance.

