## Supplemental Information for "The mechanism of coupled folding-upon-binding of an intrinsically disordered protein"

### MD simulations of the apo $N_{TAIL}$ α-MoRE

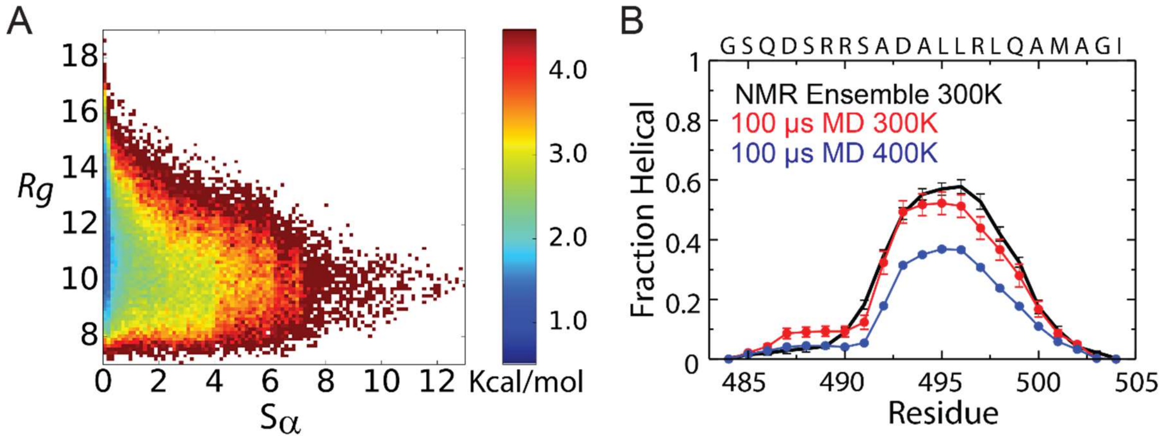


**Figure S1.** Ensemble of the apo $N_{TAIL}$ α-MoRE from a 100-μs MD simulation at 400 K. A) Free energy surface of the apo $N_{TAIL}$ α-MoRE as a function of the radius of gyration ($R_g$) and the α-helical folding order parameter Sα at 400 K. B) Comparison of the helical propensity of the apo $N_{TAIL}$ α-MoRE calculated from a 100-μs MD simulation at 400 K, a 100-μs MD simulation at 300 K, and an NMR ensemble[1] calculated with the ASTEROIDS[3] ensemble selection algorithm using NMR chemical shifts and RDCs as restraints. Helical propensities were calculated using the program STRIDE.

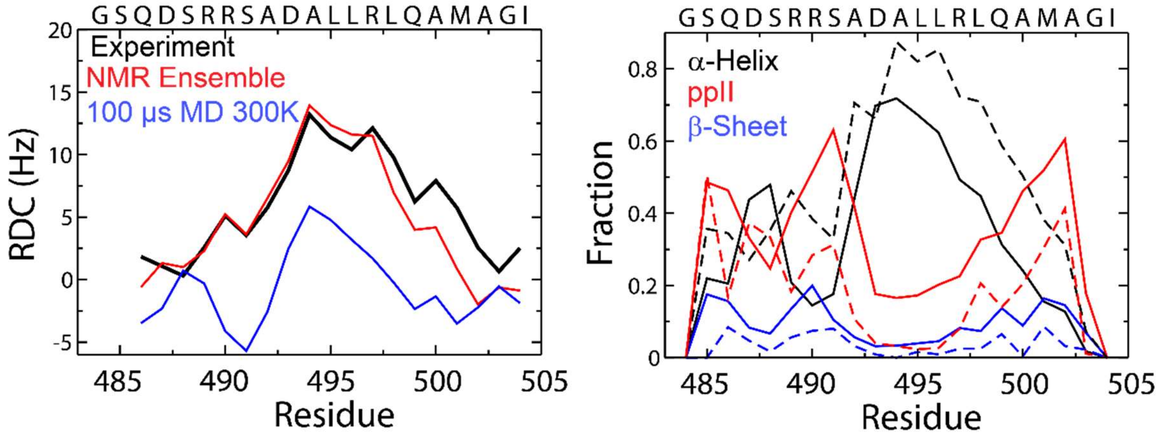


**Figure S2.** A) RDCs calculated from a 100-μs MD simulation of the apo $N_{TAIL}$ α-MoRE at 300 K (blue) and from an NMR ensemble[1] (red) are compared to experimental values (black). B) Ramachandran distribution propensities from a 100-μs MD simulation of the apo $N_{TAIL}$ α-MoRE at 300 K (solid lines) and from an NMR Ensemble[1] (dotted lines). Secondary structure propensities were calculated from previously defined Ramachandran cutoffs.[1] The disparity between the MD ensemble and experiment appears to be the result of increased populations of ppII and β sheet and decreased populations of α-helix, especially for residues 489–491.

### Agreement between NMR RDCs and 100-μs MD simulation of the apo NTAIL α-MoRE at 300 K

The finding that RDCs calculated from the MD ensemble are in worse agreement with the experimental RDCs than those calculated from the ASTERIODS ensemble appears to be in large part the result of sampling ~50% less of the helical region of Ramachandran space and ~100% more of the polyproline-II region Ramachandran space for residues R489, R490, and S491. As the helical propensities calculated by the program STRIDE, which considers both hydrogen bond geometry and backbone dihedral angles in its assignment of helical states, are similar for these residues in the MD and ASTEROIDS ensembles, this result suggests that the ASTEROIDS ensemble samples the helical region of Ramachandran space more frequently when these residues are not part of hydrogen bonded helical conformations.

### Analysis of helix and contact formation during the folding-upon-binding transition paths

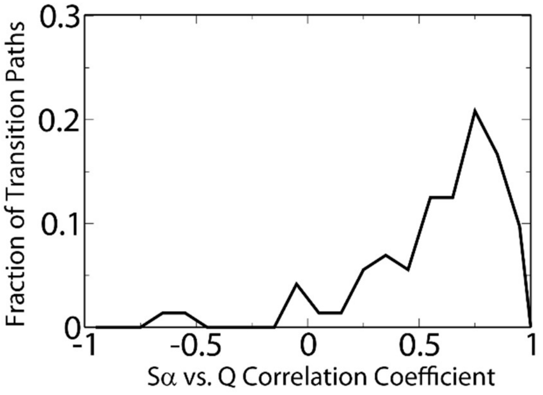


**Figure S3.** Histogram of correlation coefficients of values of Sα vs. Q for conformations sampled in 72 transition paths from an unbiased 200-μs MD simulation of the $N_{TAIL}$ α-MoRE and

XD domain run with the a99SB-*disp* force field at 400 K. The correlation coefficient of Sα vs. Q taken over all transition paths is 0.55.

| | Sα increases | Sα decreases | Sα equal |
|---|---|---|---|
| Q increases | 0.62 | 0.26 | 0.12 |
| Q decreases | 0.42 | 0.46 | 0.12 |
| Q equal | 0.33 | 0.49 | 0.18 |
| | Q increase | Q decrease | Q equal |
| Sα increases | 0.66 | 0.24 | 0.11 |
| Sα decreases | 0.42 | 0.37 | 0.21 |
| Sα equal | 0.55 | 0.26 | 0.19 |

**Table S1.** Probabilities of cooperative and noncooperative changes in Q and Sα calculated over all transition paths at 400 K. In the upper part, $p(\Delta S\alpha|\Delta Q)$ is reported; in the bottom half, $p(\Delta Q|\Delta S\alpha)$ is reported. Transitions were calculated using a 50-ns lag. Increases and decreases were determined using intervals of 0.01 for values of Q, and intervals of 0.10 for Sα. Values of Q and Sα were smoothed using a running average with a 2-ns window.

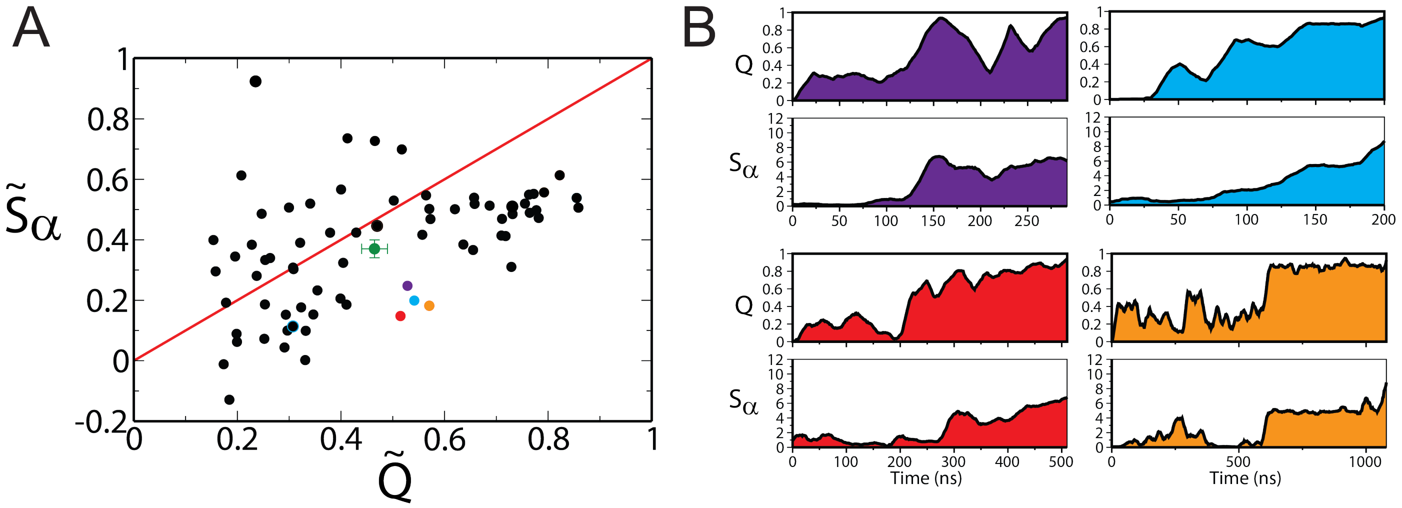


**Figure S4**. The time evolution of Sα and Q for the four representative transition paths with induced-folding character from an unbiased 200-μs MD simulation of the NTAIL α-MoRE and XD domain run with the a99SB-*disp* force field at 400 K. A) Comparison of $\tilde{S}_{\alpha}$ and $\tilde{Q}$ (normalized time averages of Sα and Q) for all observed transition paths. The four representative transition paths are shown with purple, blue, red, and orange circles. B) The time-evolution of Sα and Q for the selected transition paths.

### Influence of temperature on the folding-upon-binding mechanism

We examined if the observed mechanisms of folding-upon-binding change with temperature by running 25 unbiased binding simulations at 300 K, in which an unfolded conformation of the $N_{TAIL}$ α-MoRE was placed in a water box containing the XD domain, and the simulation continued until a stable complex formed. A further 100-μs simulation was run at 300 K in the bound state to compute the bound-state equilibrium properties. In 22 of these 25 simulations, a transition was observed in which the $N_{TAIL}$ α-MoRE bound to XD in the crystallographic pose (in the remaining three simulations the $N_{TAIL}$ α-MoRE bound in the same binding groove of XD but with the opposite orientation). On average, each simulation spent 27 μs in the unbound state, which enabled the calculation of the unbound state equilibrium properties at 300 K.

We found that the distribution of folding-upon-binding mechanisms in these 22 binding events is similar to that observed at 400 K (Figures S5 and S6), but that the transition paths tended to have even more induced-folding character at 300 K; this is consistent with experimental kinetics measurements conducted at physiological temperatures, in which separate rates for encounter complex formation and subsequent folding can be clearly resolved.[2] Similarly to what was observed at 400 K, we observed that multiple folding-upon-binding transition paths populate conformations that, on average, have less helical content than the $N_{TAIL}$ α-MoRE in apo simulations at 300 K (i.e., the $\tilde{S}_{\alpha}$ is <0).

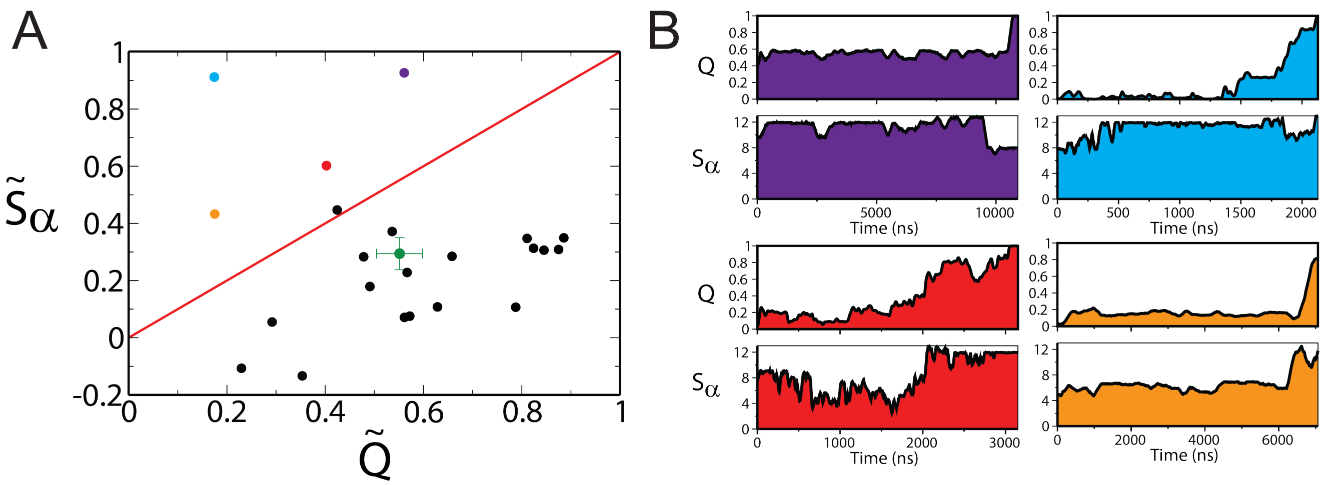


**Figure S5.** A) Comparison of $\tilde{S}_\alpha$ and $\tilde{Q}$ for binding transition paths observed in 22 unbiased MD simulations of the $N_{TAIL}$ α-MoRE and XD domain run with a99SB-*disp* force field at 300 K. Simulations were started from an unfolded conformation of the $N_{TAIL}$ α-MoRE placed in a water box with the XD domain and run until a stable native complex formed. The average value of $\tilde{S}_\alpha$ and $\tilde{Q}$ over all transition paths is shown in green. B) The time evolution of Sα and Q for four transition paths with the largest values of $\tilde{S}_\alpha$.

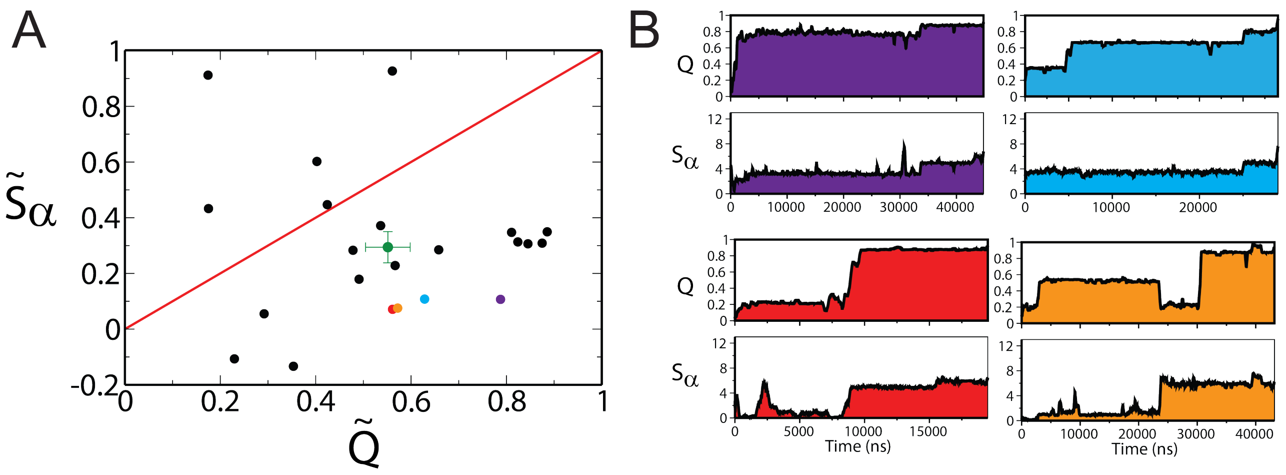


**Figure S6.** The time evolution of Sα and Q for four representative transition paths with induced folding character from unbiased MD simulations of the NTAIL α-MoRE and XD domain run with a99SB-*disp* force field at 300 K. A) Comparison of $\tilde{S}_{\alpha}$ and $\tilde{Q}$ for all observed transition paths in unbiased binding simulations. The four representative transition paths are the purple, blue, red, and orange circles. B) The time-evolution of Sα and Q for the selected transition paths.

### Optimization of the folding-upon-binding reaction coordinate

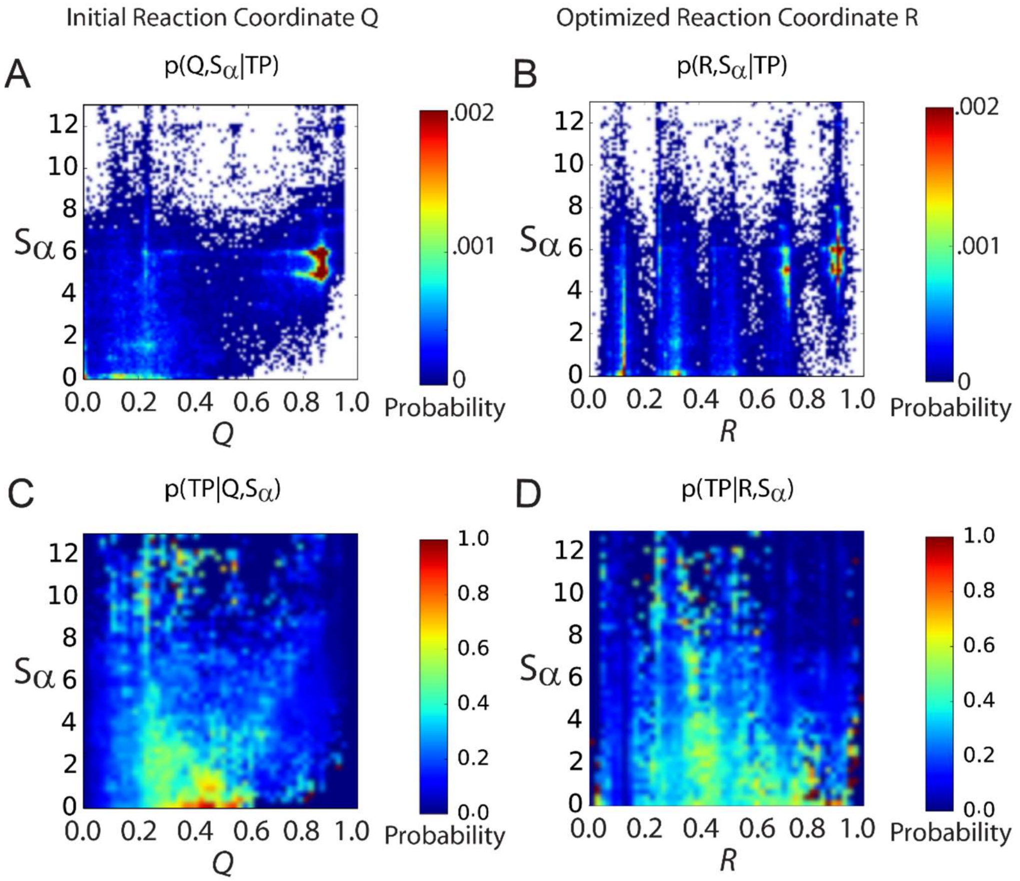


**Figure S7**. A) The probability of (Q, Sα) given that a frame is on a transition path. B) The probability of (*R,* Sα) given that a frame is on a transition path. C) The probability of being on a transition path for given values of Q and Sα. D) The probability of being on a transition path for given values of *R* and Sα.

### Calculation of a TSE

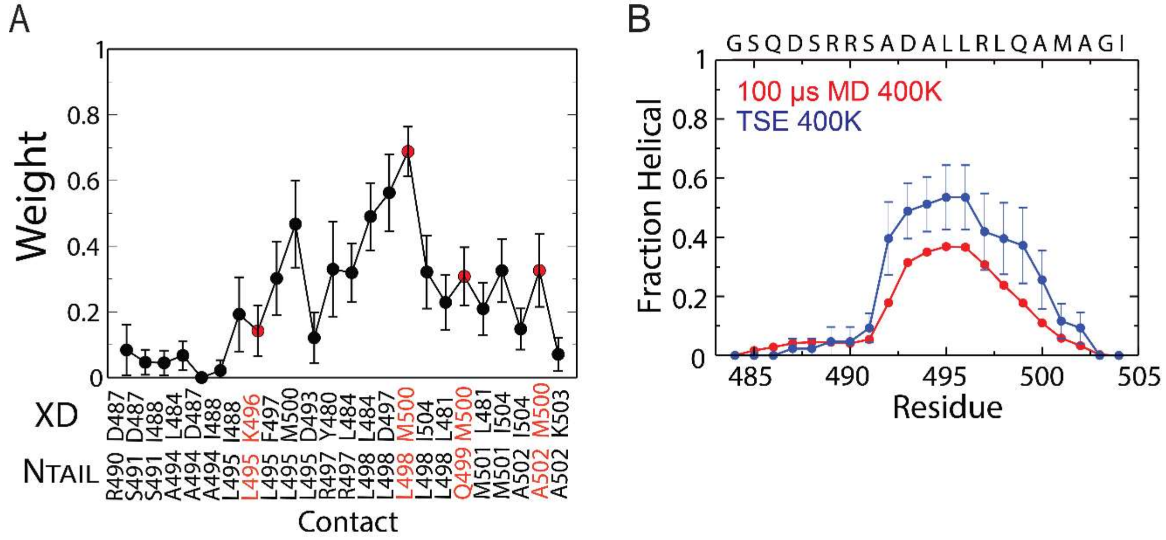


**Figure S8.** A) Average contact probability for all native intermolecular contacts in the calculated TSE. The four contacts identified as having the largest weights by the reaction coordinate optimization are colored red. B) Comparison of the helical propensity of the apo $N_{TAIL}$ α-MoRE calculated from a 100-μs MD simulation at 400 K and the TSE calculated at 400 K.

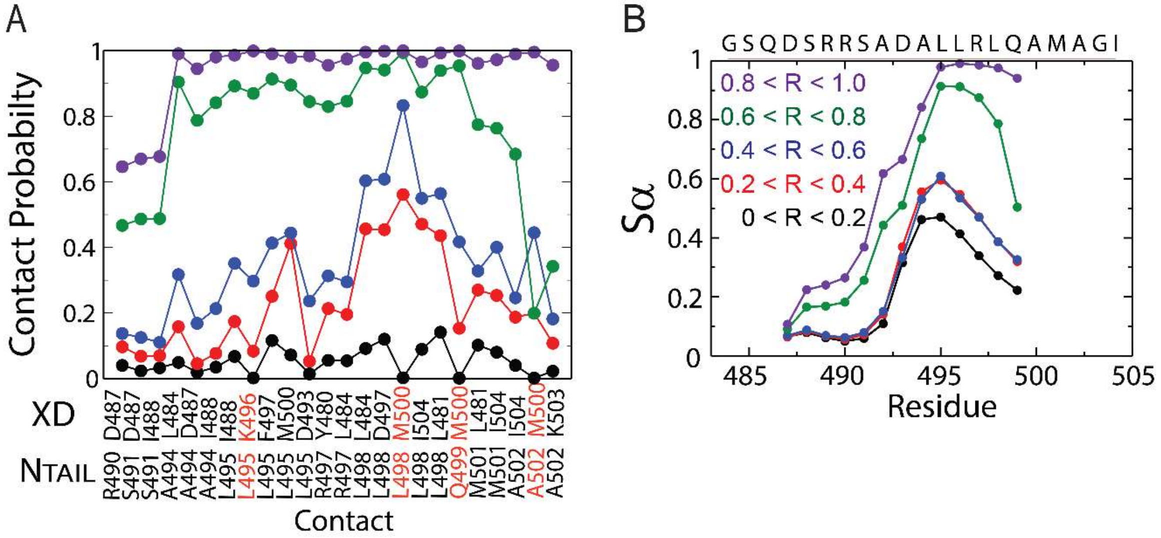


**Figure S9.** Progression of intermolecular contact and helix formation along the reaction coordinate *R*. Frames from the 200-μs 400 K trajectory were assigned into five evenly spaced bins according to the value of *R* in intervals of 0.2. A) Average contact probability for all native intermolecular contacts for each interval of *R*. B) Comparison of the Sα value for the 13 6-residue segments considered in this study. The value for each segment is plotted for the third residue of each segment (e.g., the Sα value for segment S485–R490 is plotted on residue 487).
